# Diversification models conflate likelihood and prior, and cannot be compared using conventional model-comparison tools

**DOI:** 10.1101/2021.07.12.452074

**Authors:** Michael R. May, Carl J. Rothfels

## Abstract

Time-calibrated phylogenetic trees are a tremendously powerful tool for studying evolutionary, ecological, and epidemiological phenomena. Such trees are predominantly inferred in a Bayesian framework, with the phylogeny itself treated as a parameter with a prior distribution (a “tree prior”). However, we show that the tree “parameter” consists, in part, of data, in the form of taxon samples. Treating the tree as a parameter fails to account for these data and compromises our ability to compare among models. Since accuracy of the inferred phylogeny strongly depends on how well the tree prior approximates the true diversification process that gave rise to the tree, the inability to accurately compare competing tree priors has broad implications for applications based on time-calibrated trees. We outline potential remedies to this problem, and provide guidance for researchers interested in assessing the fit of tree models.

---

Evolutionary inferences that have an explicit temporal component—when did a particular group arise, how do lineages disperse over space and diversify over time, etc.—depend on phylogenetic trees with branch lengths measured in units of time (“time-calibrated trees”). Unfortunately, character data (typically molecular sequences or morphological traits for the taxa of interest) are only informative about the amount of evolution that occurs along each branch of the tree, which is the product of the rate of evolution and the temporal duration of the branch (Felsenstein 1981; Yang 2014): a short-lived lineage that evolves quickly will experience the same amount of evolution as a long-lived lineage that evolves slowly. Consequently, methods for inferring time-calibrated phylogenies require additional information to disentangle the confounding effects of rate and time.

Methods for estimating time-calibrated trees are predominantly Bayesian (Yang 2014; Heath and Moore 2014; dos Reis et al. 2016), and disentangle rate and time through priors on rates of evolution (a “clock prior”) and on the durations of lineages (a “tree prior”). The tree prior often takes the form of a stochastic birth-death process (Kendall 1948; Yang and Rannala 1997), which is a model with parameters that govern the frequency of birth (speciation or lineage splitting), death (extinction), and sampling (*e.g*., fossilization or viral collection) events. Recently developed extensions to this class of tree model—e.g., the “serially sampled” or “fossilized” birth-death processes (Stadler 2010; Gavryushkina et al. 2014; Heath et al. 2014; Zhang et al. 2016)—also model sampling events over time, and allow researchers to estimate time-calibrated trees for mixed samples of extant and extinct lineages. Variants of these processes allow birth, death, and sampling rates to vary over time (Stadler 2011), across the branches of the tree (Maliet et al. 2019; Höhna et al. 2019), or as a function of an evolving trait or geographic region (Maddison et al. 2007; Goldberg et al. 2011). These models have the potential to reconcile major discrepancies between phylogenetic and paleontological estimates of the origin times of lineages (Ronquist et al. 2016; Marshall 2019), and allow researchers to study epidemiological dynamics as they unfold over time and space (Müller et al. 2021; Nadeau et al. 2021).

However, the power to estimate time-calibrated trees comes at a cost: because the data are unable to distinguish rate and time, estimates of time-calibrated trees are highly sensitive to the clock and tree priors (Condamine et al. 2015; Lee and Yates 2018; Wright et al. 2020; May et al. 2021), which indicates the need for careful model comparison. Comparing the fit of competing models in a Bayesian framework involves computing the marginal likelihood of each model, which is the probability of the data averaged over all parameters in proportion to their prior probability. It is therefore critical that we clearly distinguish between data (variables that are governed by likelihood functions) and parameters (variables that are governed by prior distributions) when comparing models.

Here, we argue that treating birth-death models as priors in Bayesian phylogenetic analysis is conceptually problematic: because birth-death models produce data (*i.e*., taxon samples), some of the probability of the tree should be viewed as part of the likelihood function. We demonstrate that this failure to accurately separate data from parameters compromises our ability to compare the fit of competing tree models. We quantify the magnitude of the error and provide both theoretical and practical solutions to the problem of comparing among tree models. Finally, we discuss the broader implications of this issue, especially the positive aspects of treating taxon samples as data in phylogenetic analyses.

## Background

In a standard Bayesian phylogenetic analysis under a birth-death model (Yang and Rannala 1997; Heath et al. 2014; Zhang et al. 2016), we imagine that a phylogeny, Ψ, evolves under a birth-death process with parameters *θ_Ψ_*, which consist of speciation, extinction, and sampling parameters (*λ, μ*, and *ϕ*, respectively). Additionally, we imagine that a set of characters (e.g., molecular and/or morphological traits) evolve along the branches of the tree according to a defined model with parameters *θ_x_*, giving rise to a character dataset, X.

In the traditional formulation (Yang and Rannala 1997; Zhang et al. 2016), the tree is treated as a parameter with a prior distribution, and the Bayesian model is represented as:

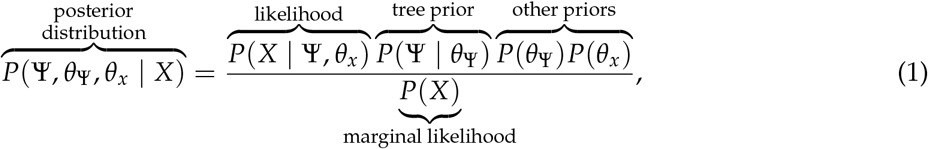

where the posterior distribution represents our belief about the parameters after observing the data, which is proportional to the probability of the data given the parameters (the likelihood) times the prior probabilities of the tree and other model parameters (our prior belief about plausible parameter values before we collected the data). The denominator is the marginal likelihood, which is the likelihood function averaged over all possible parameter values in proportion to their prior probability. The marginal likelihood is often very difficult to compute directly, so we usually approximate the posterior distribution using Markov chain Monte Carlo (MCMC;Metropolis et al. 1953), which conveniently sidesteps the need to know the marginal likelihood. However, the marginal likelihood plays a central role in Bayesian model comparison, as we describe below.

## Samples as Data

This traditional formulation, which treats the tree model as a prior distribution, is conceptually problematic. To illustrate our point, we consider a fairly simple model where diversification, character evolution, and sampling occur independently. We imagine there is a process of lineage diversification that produces the tree, as well as a process of character evolution that evolves along the branches; together, these processes produce the true (but unobserved) history of the group (Fig. 1A). The process also produces samples of extant and/or extinct lineages according to a random process governed by the sampling parameters (Fig. 1B). Finally, in order to estimate the phylogenetic relationships among the samples, a researcher may record the character data for each sample (Fig. 1C). The traditional phylogenetic treatment implicitly assumes that the character data generated in the third step are observations, and their probability is assigned to the likelihood of the model. In contrast, the samples observed in the second step are not considered observations; rather, they are considered part of the (unobserved) tree, and their probability is assigned to the prior probability of the tree model.

**Figure 1:**
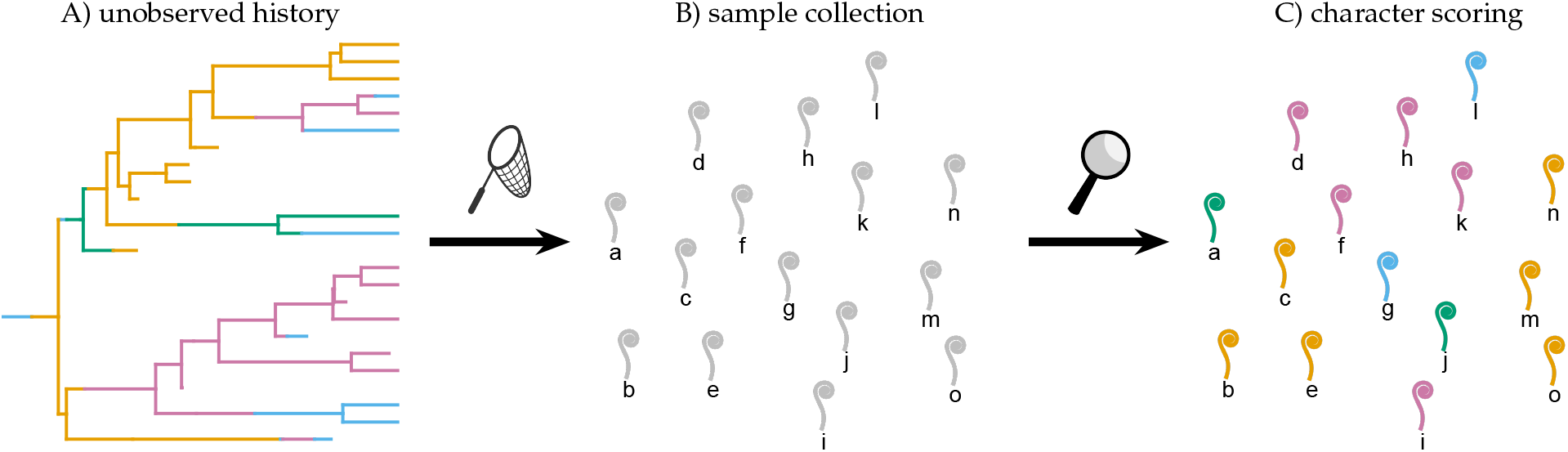
The phylogenetic process and data. A) We imagine that there is a true—but unobserved—history for our study group. This unobserved history is the outcome of random processes of lineage diversification (which gives rise to the tree topology and branch lengths) and character evolution (which assigns colors to the branches). Inferring this unobserved history—and the processes that generated it—from the samples is the focus of a phylogenetic analysis. B) We collect samples of extant and/or extinct members of our study group (grey fiddleheads). C) We score each sample for character data (color of each fiddlehead), and use those data to estimate the phylogeny of our study group. Traditionally, we treat the probability of the character data (C) evolving under a particular model of character evolution as the likelihood function, but the probability of the samples and their occurrence times (B) as part of the prior probability of the tree.

However, the samples themselves are observations that carry information about the underlying diversification process. Indeed, paleontologists regularly treat samples (fossil occurrences) as data, and use them to infer diversification rates in the absence of character data (*e.g*., Foote 2001; Silvestro et al. 2014); likewise, epidemiologists use the number of cases over time to estimate the parameters of SIR models (which share many features with birth-death models;Kühnert et al. 2016; MacPherson et al. 2021). From this perspective, it is clear that part of the “prior” probability of the tree in the traditional formulation, *P*(Ψ| *θ*_Ψ_), should in fact be treated as part of the likelihood.

Based on the fact that taxon samples are data, we present an alternative formulation of the Bayesian phylogenetic model as:

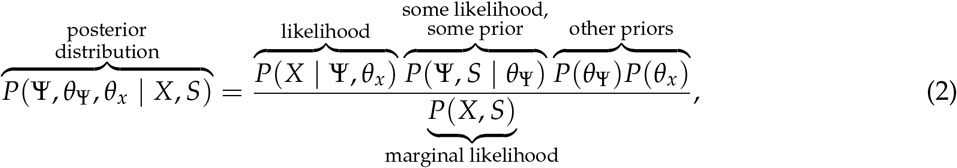

where *S* represents the taxon sample data (*e.g*., the number of samples and their ages). This representation differs from the traditional one in two main ways: first, the joint probability of the tree and the samples is a mix of likelihood and prior probability; second, the marginal likelihood is the probability of the character data *and* the samples.

Because the posterior distribution of a Bayesian model does not depend on the labeling of the likelihood and prior terms, and MCMC obviates the need to compute the marginal likelihood, the traditional and alternative formulations should produce the same posterior distributions (as we explain in section “Quantifying the Discrepancy”). In contrast, Bayesian model comparison depends on computing the marginal likelihood, so failing to treat taxon samples as data will prevent us from accurately comparing among tree models.

## Simulation Study

We simulated data to demonstrate the consequences of treating birth-death models as priors on Bayesian model comparison. The standard method for comparing Bayesian models is the Bayes factor (Jeffreys 1935; Kass and Raftery 1995). Bayes factors can be calculated in two ways: the first, and most commonly used, is to compute the marginal likelihoods of each model—using a special algorithm, e.g, the stepping-stone (SS) sampler (Xie et al. 2011)—and take the ratio:

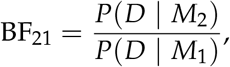

where *D* is the dataset and *P*(*D* | *M_i_*) is the marginal likelihood of model *M_i_*. Bayes factors larger than one (or zero for log BFs) indicate support for model *M*_2_; below one (zero for log BFs) indicate support for model *M*_1_. An alternative—but theoretically equivalent—approach is to place a prior on the competing models, and then estimate the corresponding posterior probability of each model, e.g., using reversible-jump (RJ) MCMC (Green 1995). We can then compute Bayes factors as:

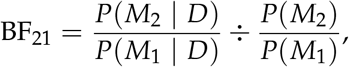

where *P*(*M_i_*) and *P*(*M_i_* | *D*) are the prior and posterior probability of model *i*, respectively.

To demonstrate that these approaches are equivalent, we used simulated data to compare two models that differ by the assumed model of molecular evolution, but use the same tree model. Marginal likelihood and RJ approaches to calculating the Bayes factor produce the same values (Fig. 2A), as expected. However, when we compare the fit of competing tree models, the two approaches disagree (Fig. 2B,C). This error is consistent with the marginal likelihoods being calculated incorrectly for tree models and is not conservative—Bayes factors computed using marginal likelihoods are not simply lower in magnitude than those computed using posterior model probabilities— and can result in Bayes factors favoring the wrong model (the model that would be disfavored if the Bayes factors were calculated correctly; Fig. 2, shaded regions). (See Supplementary Material section S1 for more details about these analyses, including the adaptive power-posterior algorithm used to calculate the marginal likelihoods.)

**Figure 2:**
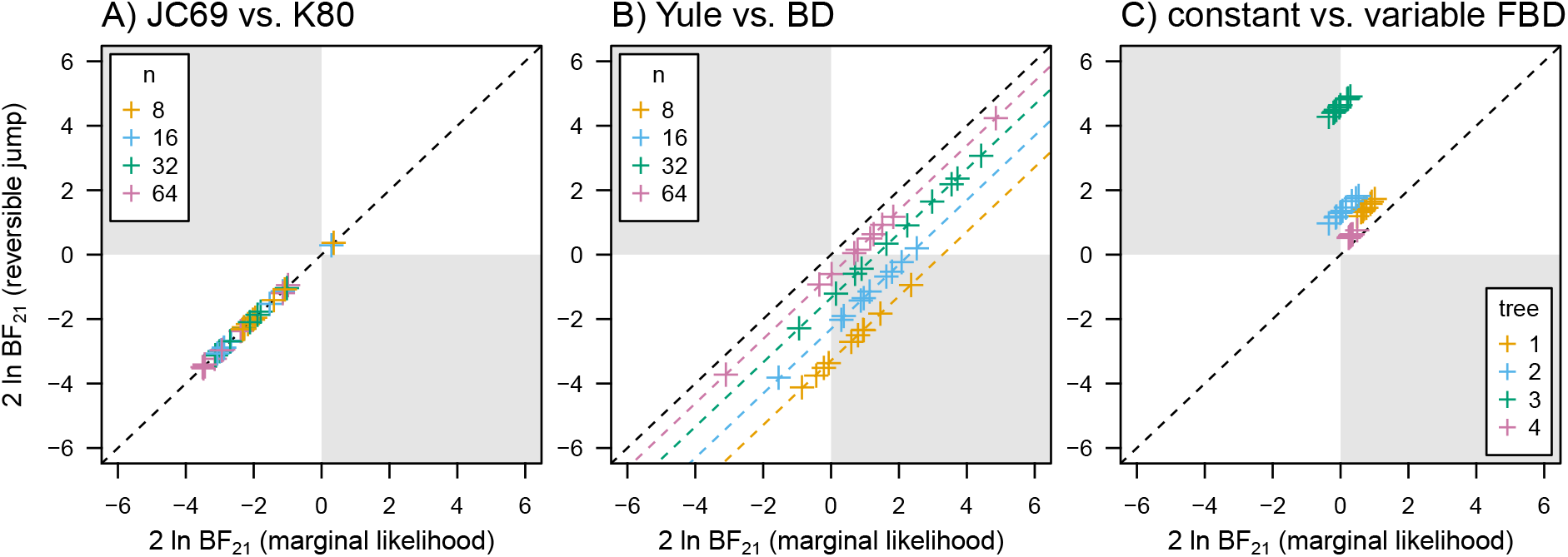
Consequences of treating birth-death models as priors on model comparison. A) The fit of two models of molecular evolution—JC69 (Jukes and Cantor 1969) (the true model) and K80 (Kimura 1980)—to simulated datasets of *n* contemporaneous samples calculated with Bayes factors (BFs) computed using a stepping-stone sampler (SS, based on marginal likelihoods;Xie et al. 2011) and reversible-jump MCMC (RJ, based on posterior model probabilities;Green 1995). On the two-log scale, positive values indicate support for the second model, and negative values indicate support for the first model. The BFs computed by these two methods are the same. B) The fit of two birth-death processes—the Yule model (with no extinction rate parameter) and the standard birth-death (BD) model (the true model)—to the datasets from panel A. There is a constant discrepancy between the two BFs that depends on the number of lineages (colored points), which is equal to the marginal likelihood ratio of the samples under the two models (dashed lines). This discrepancy can result in the two methods preferring different models (shaded regions). C) The fit of two fossilized birth-death models—one with a constant fossilization rate and another with a fossilization rate that varies over time (the true model)—to simulated datasets of non-contemporaneous samples. The resulting BFs demonstrate the same constant discrepancy as in the contemporaneous case, though in this case the ratio of the marginal likelihood of the samples is effectively impossible to calculate.

## Empirical Analysis

To explore the impact of this error on empirical inferences, we reanalyzed data from our study of marattialean ferns (May et al. 2021). Our original study reached the surprising conclusion that there was no evidence for fossilization rates varying over time, despite the fact that the majority of our fossil specimens were drawn from a narrow time interval (the Pennsylvanian). In our re-analysis, as in our original analysis, marginal likelihoods favor a model with constant fossilization rates over one that allows fossilization rates to vary (2 ln BF ≈3 in favor of the constant-rate model). However, Bayes factors calculated using RJMCMC very strongly favor the model that allows fossilization rates to peak in the Pennsylvanian (2lnBF ≈18 in favor of the variable-rate model). In other words, conventional marginal-likelihood-based BFs incorrectly indicated strong evidence for a decisively worse model. (See Supplementary Material section S2 for more details about these analyses.)

The choice of tree prior results in large differences in divergence-time estimates for this dataset; in some cases, mean node ages differ by up to ≈15 million years (Fig. 3). Additionally, the two models produce strikingly different inferences about the underlying macroevolutionary process. The variable-rate model predicts a ≈20-fold increase in the fossilization rate during the Pennsylvanian, consistent with the large number of fossils collected in that subperiod (Fig. S3). Furthermore, under the constant model, we infer that there were about 400 (95% credible interval [36,4789]) marattialean lineages during the Pennsylvanian, which drops by nearly half—to 225 (95% CI [25,2757]) lineages— under the variable model: the constant model struggles to produce the observed number of fossils in this epoch without implying a very large number of unsampled lineages (Fig. S4).

**Figure 3:**
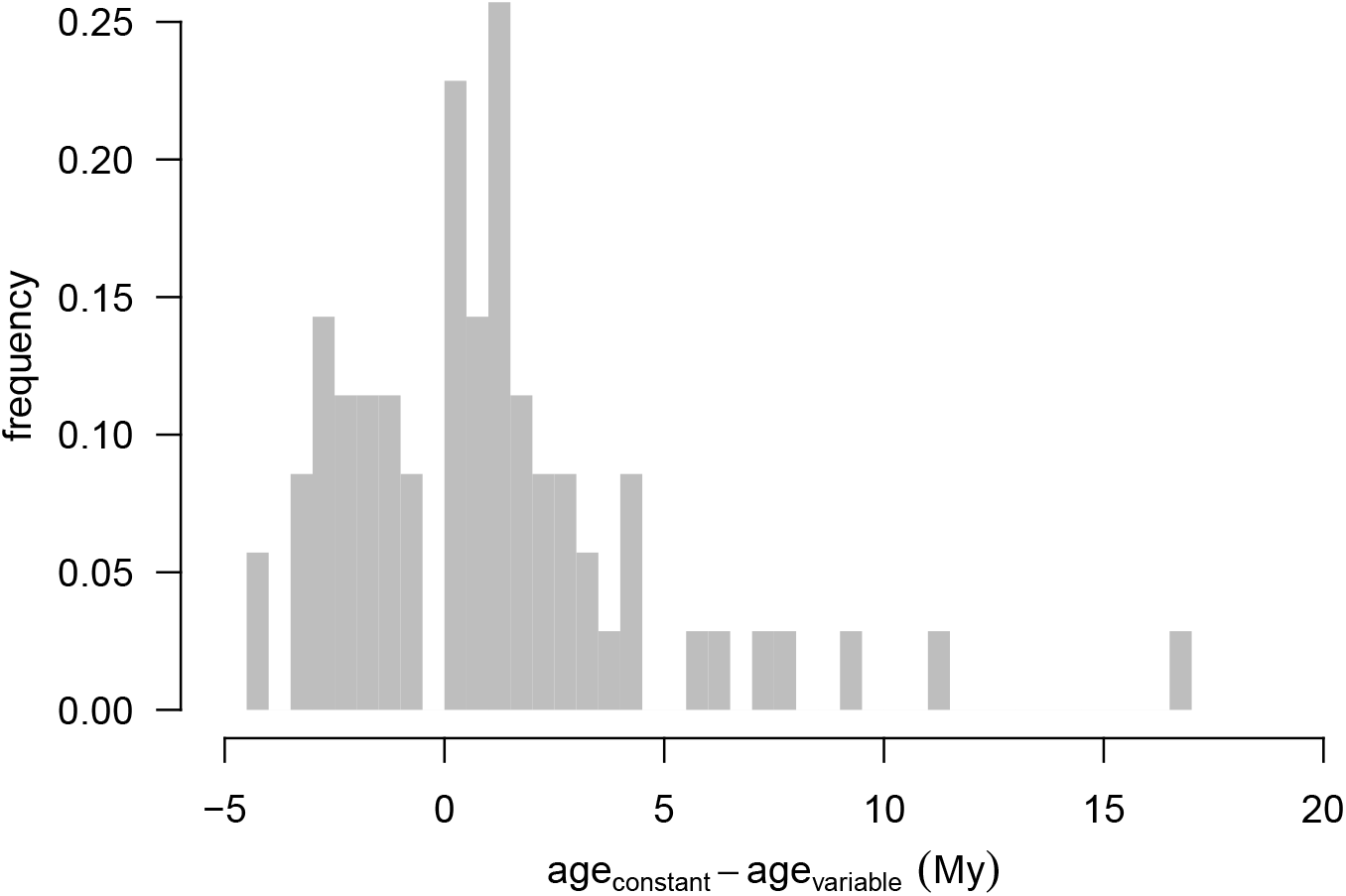
Empirical consequences of using a worse-fitting tree model. We estimated divergence times for marattialean ferns (May et al. 2021) under a model with constant fossilization rates and another model that allowed fossilization rates to vary among epochs. We plot a histogram of absolute differences in node-age estimates (computed as the posterior mean estimate of a given node age [in millions of years] under each model) under the two tree models, across all nodes in the tree. Node-age estimates disagree by up to fifteen million years under these two models, and are, on average, older under the constant-rate model than the variable-rate model.

## Quantifying the Discrepancy

We can quantify the Bayes-factor discrepancies induced by treating tree models as priors using the principles of sequential Bayesian analysis. The parameters of the prior distributions (“hyperparameters”) we choose for a Bayesian model represent our prior belief about plausible parameter values, and in principle reflect our previous experiences with analyzing relevant data (or ignorance, if we have no previous experience). In a sense, when informed by previous analysis, these hyperparameters encapsulate the information in the previous datasets about the parameters, *i.e*., they can be viewed as “old” data. When we analyze a “new” dataset, we update these “old” (prior) beliefs accordingly. We can repeat this process indefinitely, as we collect additional datasets. This sequential Bayesian updating process is the basis of Lindley’s aphorism that “today’s posterior is tomorrow’s prior” (Lindley 1972).

When we perform a Bayesian phylogenetic analysis under a birth-death model, we can imagine collecting two datasets. We first collect samples, *S* (Fig. 1B). We then infer the parameters of the birth-death model, *θ*_Ψ_, directly from this dataset. We can write the posterior distribution of this first step:

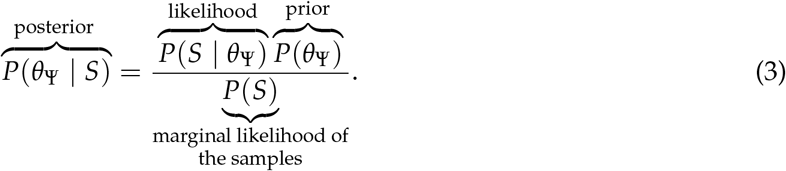

Next, we assemble a character dataset, *X*. Rather than re-doing the initial analysis, we can apply the principle of sequential Bayesian updating and use the first posterior as a prior in our second analysis:

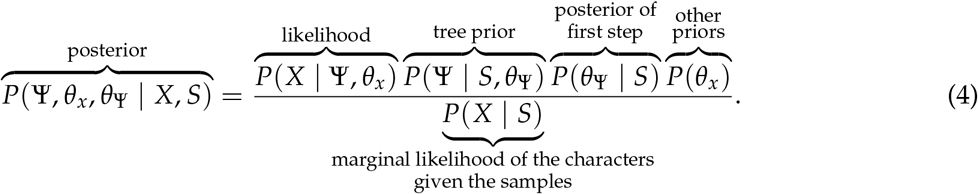

We can confirm that the posterior distribution from the sequential analysis is equivalent to the joint analysis, (2), by substituting (3) into (4) and recognizing that the marginal likelihood of the joint analysis can be factored as *P*(*X, S*) = *P*(*X* | *S*)*P*(*S*).

Importantly, the only term in the second step that is labeled as a likelihood function is the probability of the character data, which is equivalent to the traditional formulation (Eq. 1). Marginal likelihoods for the second step of a sequential analysis (or using the traditional formulation) will therefore be different from the joint analysis by the marginal likelihood of the samples, *P*(*S*), and consequently Bayes factors will be off by the ratio *P*(*S* | *M*_2_) ÷ *P*(*S* | *M*_1_). By contrast, Bayes factors computed using posterior-model probabilities from RJ MCMC should be unaffected because they are based on the posterior probabilities of the models, which do not depend on distinguishing between likelihood and prior terms. Indeed, when we calculate the marginal likelihood of the samples for data simulated under simple birth-death models with contemporaneous (e.g., extant) samples, it perfectly explains the discrepancy we observe between SS and RJ MCMC estimates of the Bayes factor (Fig. 2B, middle, dashed lines). Unfortunately, while this quantity can be computed for birth-death processes for contemporaneous samples, straightforward solutions are currently unavailable for processes that generate non-contemporaneous samples, as in Fig. 2C.

Intuitively, Bayes factors for tree models, calculated from marginal likelihoods, measure the evidence in the character data for the competing birth-death models, but ignore the evidence from the samples. By contrast, Bayes factors based on posterior model probabilities include all the relevant evidence from the samples about the relative fit of the birth-death models.

## Prognosis

Our simulations and theoretical exploration allow us to provide some advice about how to overcome the challenges of comparing tree models. The most straightforward existing solution is to use RJ MCMC for tree-model comparison. Although this approach has the additional advantage of averaging estimates of the tree (and all other parameters) over the uncertainty in the underlying tree model, these algorithms need to be tailored to specific sets of models, which constrains the types of models we can compare.

A more satisfying, generic solution would be to develop the mathematical machinery to separate the probability of the samples from the probability of the tree. In principle, we can refactor the joint probability of the tree and samples into unambiguous likelihood and prior components:

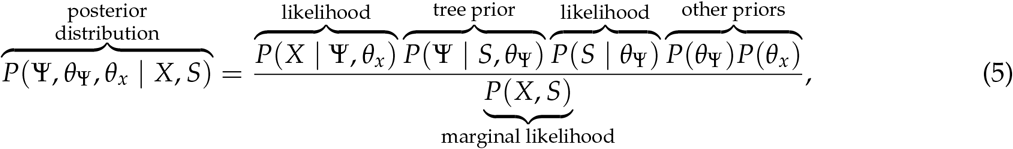

in which case methods for computing marginal likelihoods that depend on labeling likelihoods would work as intended.

While the necessary quantities *P*(Ψ| *S*, *θ*_Ψ_) and *P*(*S* | *θ*_Ψ_) are relatively straightforward to compute for simple birth-death models that generate contemporaneous samples (as we demonstrate in Supplementary Material, section S3), they are seemingly intractable for models that generate non-contemporaneous samples, especially when the exact ages of the samples are uncertain (as is often the case with fossils). Additionally, more complex models where the underlying history of the group depends on the sampling process, such as epidemiological models where sampling events correspond to extinction, or cases when the sampling rate depends on the state of an evolving trait as in state-dependent birth-death models (Maddison et al. 2007), may require different factorizations that the one presented above to separate likelihoods from priors, and will almost certainly be similarly intractable.

Methods for inferring diversification rates from fossil data (Silvestro et al. 2014; equation 7 from Stadler et al. 2018) assume the birth and death times (*b* and *d*) of every lineage are known in order to calculate the joint probability *P*(*S*, *b*, *d* | *θ*_Ψ_), and then use MCMC to numerically average over the unknown birth and death times. The strength of this approach is that it sidesteps the need to compute *P*(*S* | *θ*_Ψ_), but the tradeoff is that it does not allow us to compute *P*(*S* | *θ*_Ψ_), and therefore cannot be used to compare birth-death models.

Thinking of taxon samples as data has additional positive consequences, beyond allowing for accurate model comparison. For example, it provides a rigorous basis for using posterior-predictive simulation (PPS;Gelman et al. 1996; Bollback 2002; Brown 2014) to assess the absolute fit (model adequacy) of birth-death models (see Supplementary Material, section S4). In contrast to Bayes factors, which compare the relative fit of competing models, PSS evaluates how well a given model describes the true data-generating process by comparing simulated data (e.g., the number and ages of taxon samples) to the corresponding observed data. As an absolute measure of model fit (as opposed to the relative measure of fit provided by Bayes factors), PPS helps not only to establish confidence (or skepticism) in particular estimates, but also to identify specific weaknesses in existing models, which enables further model development.

More abstractly, treating taxon samples as data actually justifies comparing among tree models with Bayes factors in the first place. While marginal likelihoods are sensitive to different prior choices, Bayesian model comparison is not intended as a tool for comparing the fit of prior distributions. Indeed, the justification for a given prior rests in how well it characterizes a researcher’s belief about plausible parameter values before seeing the data, so assessing its quality based on its fit to a new dataset is philosophically questionable. By contrast, comparing the fit of birth-death models to observed taxon samples is a perfectly reasonable—indeed, the intended—application of model comparison.

## Data availability

Supplementary scripts and data can be found in the Zenodo repository http://doi.org/10.5281/zenodo.5072533 and the GitHub repository https://github.com/mikeryanmay/bd_bayes_factors/releases/tag/initial_submission.

## Acknowledgments

We thank members of the Rothfels lab, Brian Moore, Carrie M. Tribble, Ixchel González Ramírez, Jenna T. B. Ekwealor, Ammon Thompson, Jiansi Gao, and the editor and X anonymous reviewers for providing valuable feedback on this manuscript. This research was supported by the National Science Foundation (NSF) grant DEB-1754705 to CJR and Cindy V. Looy. This research used the Savio computational cluster resource provided by the Berkeley Research Computing program at the University of California, Berkeley (supported by the UC Berkeley Chancellor, Vice Chancellor for Research, and Chief Information Officer).

