## Supplemental Material for "Diversification models conflate likelihood and prior, and cannot be compared using conventional model-comparison tools"

#### Contents

**S1 Simulation Study** **S2**
S1.1 Marginal likelihood estimation . . . . . S2
S1.2 Simulation 1: Molecular substitution models . . . . . S2
S1.3 Simulation 2: Contemporaneous birth-death processes . . . . . S3
S1.4 Simulation 3: Fossilized birth-death processes . . . . . S3
S1.5 Results . . . . . S3

**S2 Empirical Analysis** **S5**
S2.1 Analysis . . . . . S5
S2.2 Results . . . . . S6

**S3 Factorizing Bayes' Theorem** **S9**

**S4 Posterior-Predictive Simulation with Samples as Data** **S11**

### S1 Simulation Study

To demonstrate the issues that arise from not treating samples as data, we performed a series of experiments in `RevBayes` (Höhna et al. 2016), which we selected for convenience; the problems we demonstrate are inherent to the standard treatment of tree models as priors and are independent of any particular implementation of these models. For each of the three experiments below, we performed a series of simulations under a specific model, and then compared the fit of a pair of competing models to these simulated data, using Bayes factors calculated with marginal-likelihood- and posterior-probability-based approaches. For the marginal-likelihood-based approach, we computed Bayes factors using a power-posterior algorithm, stepping-stone MCMC (SS MCMC; Xie et al. 2011), and for the posterior-probability-based approach we used reversible-jump MCMC (RJ MCMC; Green 1995). In all cases, we fixed the tree topology to the true value for computational tractability, but estimated the node ages, the birth-death parameters, and the parameters that governed the process of character evolution; for fossilized birth-death datasets, we also estimated whether each fossil was a sampled ancestor (*i.e.*, whether it was sampled along a branch leading to another sample).

#### S1.1 Marginal likelihood estimation

Initial experiments indicated that sufficiently precise marginal-likelihood estimates using SS MCMC would be computationally prohibitive for large trees and sequence datasets. We therefore simulated relatively small trees and character datasets (exact sizes described below). Further, we implemented an adaptive power-posterior algorithm in `RevBayes`, similar to the one proposed by Friel et al. (2014). Power-posterior algorithms work by running a set of Markov chains, each with a "power", $\beta_i$, ranging between 0 and 1. For any given $\beta_i$ (a "stone"), the chain samples from the distorted posterior distribution:

$$P_i(\theta \mid X) \propto P_i(X \mid \theta)^{\beta_i} P(\theta),$$

so that $\beta = 1$ corresponds to sampling from the posterior, and $\beta = 0$ corresponds to sampling from the prior. The sampled likelihood values among the separate stones—$P_i(X \mid \theta)$—can then be used to estimate the marginal likelihood, *e.g.*, using the stepping-stone estimator (Xie et al. 2011). Usually, the number of stones and the values of $\beta$ are fixed in advance, but in our analyses we found that accurate marginal-likelihood estimates demanded a large number of stones, so adopted an adaptive approach. Briefly, our adaptive algorithm begins with two stones, $\beta_1 = 1$ and $\beta_2 = 0$, and then places additional stones until the estimate of the marginal likelihood converges; as with the original algorithm, the number of MCMC samples per stone is fixed in advance.

For each analysis described below, we performed two replicates to ensure stability of marginal-likelihood and posterior-ratio estimates. Our simulated data and code (including specific parameter settings for simulations and analyses) are available at Zenodo (`http://doi.org/10.5281/zenodo.5072533`) and GitHub ( `https://github.com/mikeryanmay/bd_bayes_factors/releases/tag/initial_submission`).

#### S1.2 Simulation 1: Molecular substitution models

To demonstrate that SS and RJ MCMC compute the same BFs (and also to demonstrate that both of these methods are implemented correctly, *i.e.*, that our results are not a consequence of programming errors), we compared the fit of competing substitution models to simulated molecular datasets. We simulated ten trees under a birth-death (BD) model for each of four numbers of extant samples, $n = \{8, 16, 32, 64\}$. We assumed the tree began with two species at time $t = 1$, diversified at rates

$\lambda = 4$ and $\mu = 2$, and that all extant species were sampled. For each tree, we simulated a nucleotide dataset with 100 sites under a Jukes-Cantor (JC69; Jukes and Cantor 1969) model with rate parameter $r$ (scaled such that the expected number of substitutions per site was three). For each of the 40 simulated datasets, we computed Bayes factors between JC69 and K80 (Kimura 1980) substitution models using SS and RJ MCMC as described above, assuming the tree evolved under the true birth-death model (Fig. 2A, main text). Positive values of $2 \ln \mathrm{BF}$ indicate support for the K80 model.

#### S1.3 Simulation 2: Contemporaneous birth-death processes

Our second experiment considers the case of comparing two birth-death models for extant (contemporaneous) samples. We analyzed the same datasets simulated in the previous section, but in this case compared two tree models. The first model, $M_1$, is the Yule model (with speciation rate $\lambda$, and no extinction rate parameter), and the second model, $M_2$, is a BD model (with speciation rate $\lambda$ and extinction rate $\mu$); $M_2$ is the same tree model used to simulate the data, as described above. We computed Bayes factors between Yule ($M_1$) and BD ($M_2$) processes, again using both SS and RJ MCMC, assuming the sequence data evolved under the true JC69 substitution model (Fig. 2B, main text). In this case, positive values of $2 \ln \mathrm{BF}$ indicate support for the true model.

#### S1.4 Simulation 3: Fossilized birth-death processes

Our third experiment considers the more complex case of comparing two birth-death models for non-contemporaneous samples. We simulated four fossilized birth-death trees under a model that allowed the fossilization rate to vary. Specifically, each tree began with one lineage at time $t = 1$ (in the past, with $t = 0$ the present) and initially evolved under a fossilized birth-death model with $\lambda = 4$, $\mu = 2$, $\phi = 3$; at time $t = 0.5$ in the past, the fossilization rate changed to $\phi = 0.5$ (*i.e.*, the fossilization rate was high in the early part of the process, and decreased in the second half by a factor of six). We then simulated stratigraphic uncertainty by dividing time into 20 equally sized bins and using the boundaries of the bin that a given fossil sample fell into as the minimum and maximum ages of the sample (we assigned extant samples minimum and maximum ages of zero). For each tree, we simulated 100 binary characters under an Mk model (Lewis 2001) with rate $r$ (scaled such that the expected number of substitutions per site was three). We then compared the fit of two competing fossilized birth-death models to the simulated data. The first model, $M_1$, has constant speciation, extinction, and fossilization rates ($\lambda$, $\mu$, and $\phi$, respectively). The second model, $M_2$, is the same as $M_1$, but allows the fossilization rate to vary over time. Specifically, the initial fossilization rate (at time $t = 1$ in the past) is $\phi_1$, and at time $t = 0.5$ units in the past, it changes to rate $\phi_2$, which persists until the present ($t = 0$). $M_2$ is similar to the simulating process in that the fossilization rate is not constant, and the time of the rate change is fixed; however, for both $M_1$ and $M_2$, we assume that the speciation, extinction, and fossilization rates are unknown. For both models, we also assume that all extant species are sampled, $\rho = 1$. We computed the Bayes factors between $M_1$ and $M_2$ using SS and RJ MCMC, assuming the morphological data evolves under the Mk model (Fig. 2C, main text). Positive values of $2 \ln \mathrm{BF}$ indicate support for the variable-rate model.

#### S1.5 Results

Our results indicate that SS and RJ MCMC provide essentially identical Bayes factors when comparing models of molecular evolution (simulation 1, Fig. 2A, main text), as expected based on the theoretical equivalence of these estimators. However, these method produce disparate estimates when comparing tree models, either for contemporaneous lineages (simulation 2, Fig. 2B, main text) or when including non-contemporaneous lineages (simulation 3, Fig. 2C, main text). The discrepancy is not a consequence of a programming bug: the comparison of substitution models demonstrates that both algorithms are correctly implemented. Likewise, it is not a consequence of numerical MCMC errors: we performed two replicates of each of the analysis to confirm that Bayes-factor estimates were sufficiently precise both for SS-based estimates (Fig. S1) and RJ-based estimates (Fig. S2).

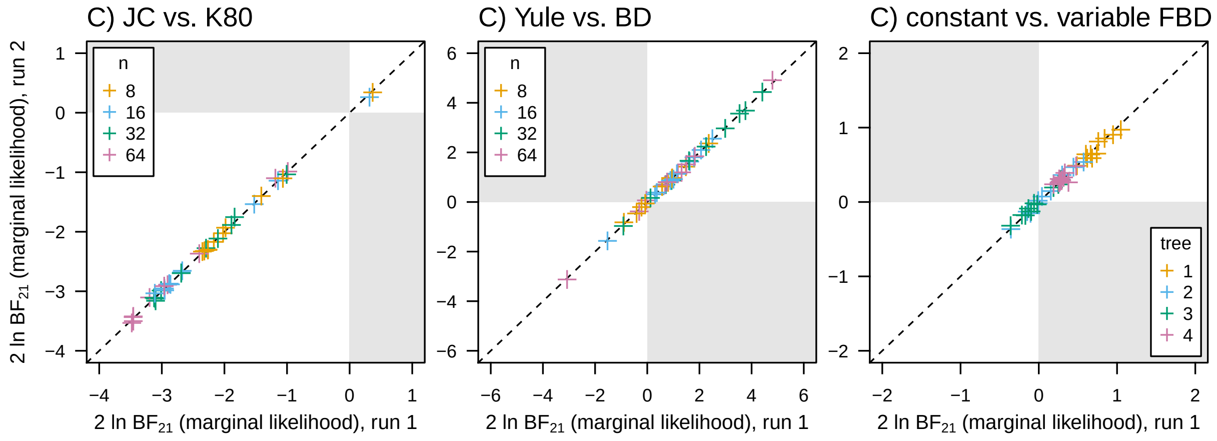


**Figure S1:** Precision of Bayes factor calculation using marginal likelihoods. Each analysis was performed twice, and the value from one run (x-axis) is plotted against the second run (y-axis). A) Bayes factors between JC69 and K80 models. B) Bayes factors between Yule and birth-death models. C) Bayes factors between a model with constant fossilization rates and one with variable fossilization rates (see caption of Fig. 1).

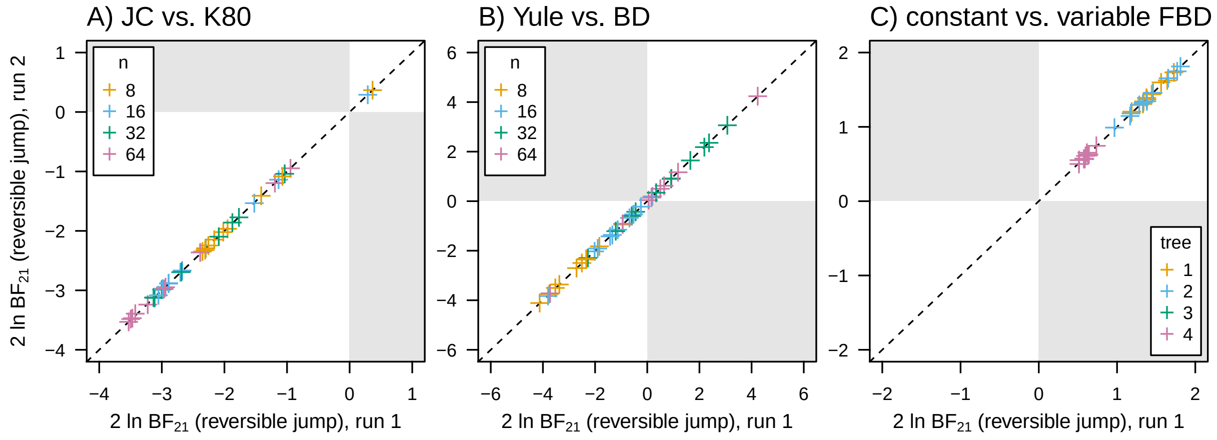


**Figure S2:** Precision of Bayes factor calculation using reversible-jump MCMC. Each analysis was performed twice, and the value from one run (x-axis) is plotted against the second run (y-axis). A) Bayes factors between JC69 and K80 models. B) Bayes factors between Yule and birth-death models. C) Bayes factors between a model with constant fossilization rates and one with variable fossilization rates (see caption of Fig. 1).

### S2 Empirical Analysis

#### S2.1 Analysis

We re-analyzed the empirical dataset of marattialean ferns from our previous study (May et al. 2021) to demonstrate the impact of not treating samples as data in a realistic model-comparison scenario, as well as to provide an example of the impact of the tree model on estimates of divergence times. For the sake of computational tractability, we included only ingroup taxa—comprising 26 extant and 45 extinct samples—and only analyzed the binary morphological data from that study (*i.e.*, we excluded the molecular data and multistate morphological characters).

We analyzed this dataset under an Mkv model (Lewis 2001) with gamma-distributed rate variation among characters (Yang 1994), and assumed that rates of morphological evolution varied across branches of the tree according to an uncorrelated-lognormal relaxed-clock model (UCLN; Drummond et al. 2006). We compared the fit of two competing birth-death models: 1) a fossilized birth-death model where speciation and extinction rates varied over time, but the fossilization rate was constant over time (the "constant" model), and; 2) a fossilized birth-death model where fossilization, speciation, and extinction rates each varied over time (the "variable" model). Specifying arbitrary variable-rate fossilized birth-death models that are amenable to efficient reversible-jump MCMC is non-trivial. We therefore used the results from our previous study to constrain the rate variation so that it was both appropriate for the dataset, and possible to specify in the existing reversible-jump machinery available in `RevBayes`. For the speciation- and extinction-rate variation, we assumed that these rates varied according to a piecewise-constant model defined by five time intervals intended to capture the major patterns present in our prior analyses: $(\infty, 323.2]$, $(323.2, 298.9]$, $(298.9, 66.0]$, $(66.0, 5.3]$, and $(5.3, 0.0]$. Within each time interval, speciation and extinction rates were drawn independently from a shared prior distribution. For the model that allowed the fossilization rate to vary, we assumed a piecewise constant model with four time intervals: $(\infty, 323.2]$, $(323.2, 298.9]$, $(298.9, 66.0]$, and $(66.0, 0.0]$, with corresponding fossilization rates $\{\psi_1, \psi_2, \psi_3, \psi_4\}$. We assumed that the rates of the first and fourth interval were the same ($\psi_1 = \psi_4$), but allowed the fossilization rates for the second and third intervals to be different, reflecting an apparent peak in fossilization rates in the Pennsylvanian (the second interval), followed by moderate fossilization rates from the Permian to the end of the Cretaceous (the third interval).

For both tree models, we fixed the tree topology to the maximum-clade-credibility (MCC) tree topology inferred in our previous study, but estimated the node ages, the fossilized birth-death parameters, the character-evolution parameters, and also whether each fossil was a sampled ancestor. We then computed the Bayes factors between the constant and variable models using SS and RJ MCMC.

#### S2.2 Results

Under the variable model, we infer extreme variation in fossilization rates: rates are inferred to be substantially higher during the Pennsylvanian than in the other time intervals, and rates from the Permian to the Late Cretaceous are also elevated compared to the first and fourth intervals (Fig. S3). This result is unsurprising, given that 24 of the 45 fossil samples come from the 23 My window that constitutes the Pennsylvanian subperiod. Despite this evident rate variation, BFs based on marginal likelihoods favor the constant model (2 ln BF $\approx$ 3, Tables S.1 and S.2). By contrast, BFs based on posterior model probabilities decisively favor the variable model (2 ln BF $\approx$ 18, Table S.2). In other words, conventional marginal-likelihood-based BFs incorrectly indicate strong evidence for a decisively worse model.

The different models produce significantly different inferences about the history of diversity for the group. We simulated 10,000 histories of lineage diversification under each model, and discretized time into many (10,000) small time intervals. We then computed the median number of lineages alive in each time interval over the group's history (Fig. S4). Under $M_1$, we predict an average peak diversity of $\approx$ 945 lineages in the Pennsylvanian; by contrast, $M_2$ predicts $\approx$ 546 lineages at that time. This discrepancy likely reflects the fact that $M_1$ requires a larger number of lineages during the Pennsylvanian in order to preserve the observed number of samples, given that fossilization rates are lower at that time relative to $M_2$ (Fig. S3).

Beside providing qualitatively different inferences about the nature of the fossilization process and the underlying history of diversification, the two tree models strongly influence divergence-time estimates for this dataset. Divergence-time estimates for young nodes are systematically more recent under the constant model, *i.e.*, the younger nodes are disproportionately pulled toward the present (Fig. S5). Presumably, this pattern reflects the fact that the fossilization rates in the Cenozoic are higher under the constant model than the variable model (Fig. S3): when fossilization rates are high, older clades in these time intervals imply more missing fossils, and are therefore "penalized" by the model (*i.e.*, younger clades are preferred).

**Table S.1: Marginal likelihoods for the constant model computed with stepping-stone sampling.** We performed four independent runs to ensure precise marginal likelihood estimates (runs 1 through 4); we report the mean and standard error of the mean (final column).

| Model | Run 1 | Run 2 | Run 3 | Run 4 | Mean (± SD) |
|---|---|---|---|---|---|
| Constant | −883.0418 | −882.8819 | −882.9806 | −882.9524 | −882.9602 (±0.03634) |
| Variable | −884.5965 | −884.5084 | −884.4403 | −884.4645 | −884.5025 (±0.03136) |

**Table S.2: 2 ln Bayes factors between constant and variable models, computed using two methods.** We performed four independent runs to ensure precise Bayes factor estimates (runs 1 through 4); we report the mean and standard error of the mean (final column).

| Method | Run 1 | Run 2 | Run 3 | Run 4 | Mean (± SD) |
|---|---|---|---|---|---|
| SS | −3.0770 | −3.3178 | −2.9194 | −3.0242 | −3.0846 (±0.0843) |
| RJ | 18.1055 | 17.9870 | 18.0041 | 18.0804 | 18.0442 (±0.0287) |

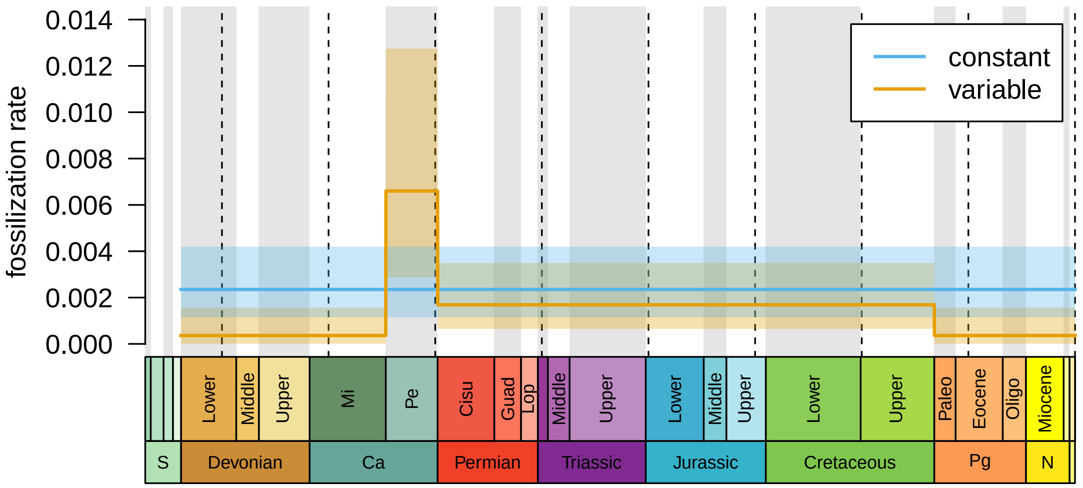


**Figure S3:** Estimated fossilization rates under the constant fossilization-rate model (blue) and the variable fossilization-rate model (orange). Dark lines correspond to the mean posterior rate at each time point, and colored regions correspond to the 95% credible interval. Dashed lines are placed at 50 My intervals.

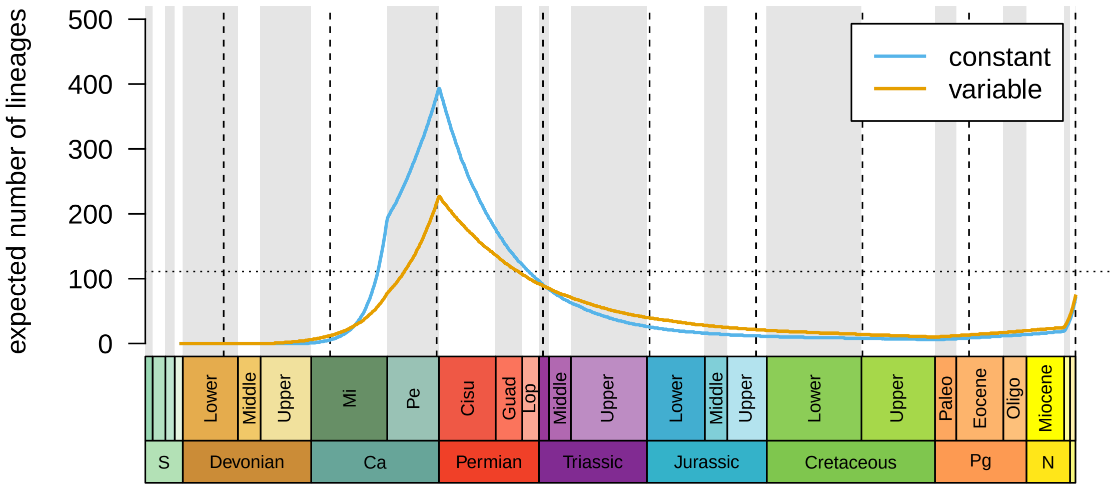


**Figure S4:** Median number of lineages over time, as predicted by the constant fossilization-rate model (blue) and the variable fossilization-rate model (orange). Vertical dashed lines are placed at 50 My intervals; horizontal dashed line is the number of extant lineages.

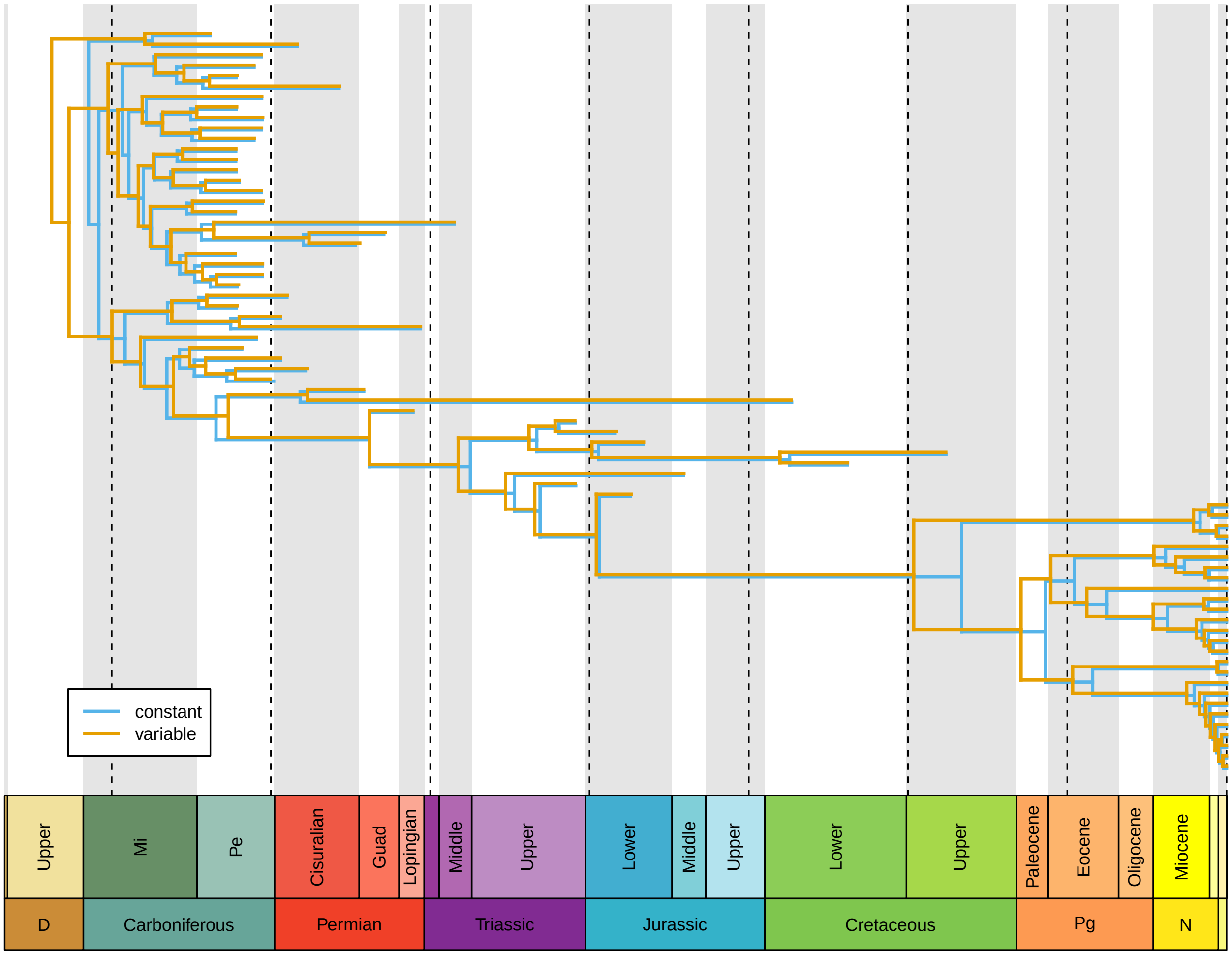


**Figure S5:** Phylogenies estimated under the constant (blue) and variable (orange) fossilized birth-death model. Node ages correspond to the the posterior mean age inferred under the model. Dashed lines are placed at 50 My intervals.

#### S3 Factorizing Bayes' Theorem

The posterior distribution of our alternative formulation for the Bayesian phylogenetic model is:

$$\overbrace{P(\Psi,\theta_\Psi,\theta_x \mid X,S)}^{\substack{\text{posterior}\\ \text{distribution}}} = \frac{\overbrace{P(X \mid \Psi,\theta_x)}^{\text{likelihood}}\overbrace{P(\Psi \mid S,\theta_\Psi)}^{\text{tree prior}}\overbrace{P(S \mid \theta_\Psi)}^{\text{likelihood}}\overbrace{P(\theta_\Psi)P(\theta_x)}^{\text{other priors}}}{\underbrace{P(X,S)}_{\text{marginal likelihood}}}, \tag{S.1}$$

such that the likelihood terms are unambiguously labeled. However, the quantities $P(\Psi \mid S,\theta_\Psi)$ and $P(S \mid \theta_\Psi)$ are not necessarily trivial to compute.

We are able to compute these quantities for Yule and birth-death models producing contemporaneous samples. The joint probability of the tree and samples can be factored as:

$$P(\Psi,S \mid \theta_\Psi) = P(\Psi \mid S,\theta_\Psi)P(S \mid \theta_\Psi), \tag{S.2}$$

where $P(S \mid \theta_\Psi)$ is the probablity of the observations $S$ (a likelihood), and $P(\Psi \mid S,\theta_\Psi)$ is the probability of the unobserved tree given the samples. Because the samples all have the same age, the probability of $S$ is just the probability of the nubmber of samples in $S$, $n$, after time $t$, which has a known analytical solution (*e.g.*, equations [20] from Höhna 2015, if the process begins with $n_t = 2$ species at time $t$):

$$\begin{aligned} P(S \mid \theta_\Psi) = P(n \mid n_t = 2,\theta_\Psi) = \\ (n-1)\left[P(n_0 > 0 \mid n_t = 1)^2 e^{-(\lambda-\mu)t}\right]^2 \left[1 - P(n_0 = 0 \mid n_t)e^{-(\lambda-\mu)t}\right]^{n-2} \end{aligned} \tag{S.3}$$

where $n_0$ is the (random) number of species at the present, $t = 0$, and assuming rates of speciation and extinction are constant for simplicity. The probability of at least one species at the present is ($n_0 > 0$):

$$P(n_0 > 0 \mid n_t = 1) = \frac{1}{1 + \int_0^t \mu e^{-(\lambda-\mu)(t-s)} ds}$$

and of zero species:

$$P(n_0 = 0 \mid n_t = 1) = 1 - P(n_0 > 0 \mid n_t = 1)$$

(*e.g.*, eq [7] from Höhna 2015). In general, the probability of $n_0 = k$ species is (eq [7] from Höhna 2015):

$$P(n_0 = k \mid n_t = 1) = \left[1 - P(n_0 = 0 \mid n_t)e^{-(\lambda-\mu)t}\right]^{k-1} P(n_0 = 0 \mid n_t)^2 e^{-(\lambda-\mu)t}.$$

The probability of the tree conditional on the number of samples also has an analytical solution (eq [21] from Höhna 2015):

$$P(\Psi \mid n,\theta_\Psi) = \frac{1}{n-1}\prod_{i=2}^{n-1}\left[\frac{i\lambda P(n_0 = 1 \mid n_{t_i} = 1)}{1 - P(n_0 > 0 \mid n_t = 1)e^{-(\lambda-\mu)t}}\right], \tag{S.4}$$

where $t_i$ is the age of the $i^{\text{th}}$ node in the tree (with the root indexed $i = 1$ and the remaining internal nodes indexed arbitrarily). Effectively identical equations are available both in Yang and Rannala (1997) and Gernhard (2008).

Together, equations (S.3) and (S.4) allow us to compute (S.2). We re-analyzed our simulated data using this formulation, demonstrating that it provides correct marginal-likelihood estimates (Fig. S6).

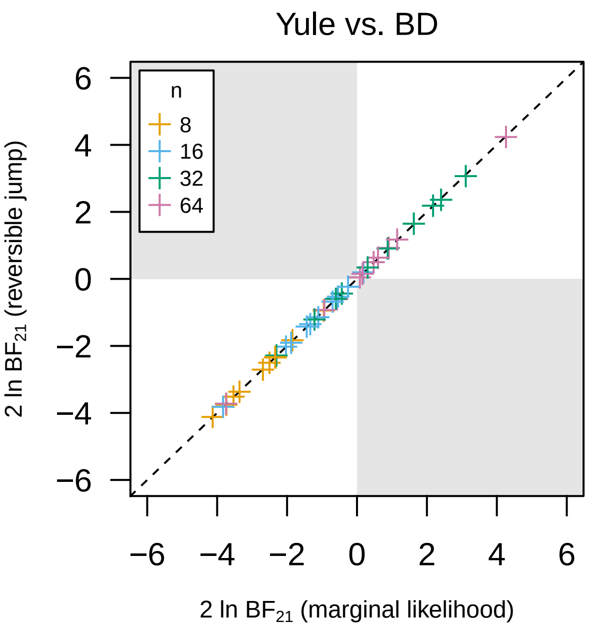


**Figure S6: Bayes factor discrepancies are resolved by refactoring Bayes' theorem.** We compared the fit of two birth-death processes—the Yule model (with no extinction rate parameter) and the standard birth-death (BD) model—to datasets simulated under the BD model. We corrected the likelihood function according to equation (S.1). As expected, there is no discrepancy between the BFs calculated using these corrected marginal likelihoods and reversible-jump MCMC.

#### S4 Posterior-Predictive Simulation with Samples as Data

While Bayes factors are useful for comparing the *relative* fit of competing models, they provide no guarantee that the best model adequately describes the process that gave rise to the observed data. Posterior-predictive simulation (PPS; Gelman et al. 1996) is a Bayesian tool that fills this gap by assessing model adequacy—whether our inference model provides an adequate description of the true process that produced our observed dataset—and is therefore useful for assessing *absolute* model fit. Generally, the procedure works by drawing parameters of the model from their joint posterior distribution (*e.g.*, as produced by an MCMC analysis), simulating new datasets under these parameters, and checking whether the simulated data resembles the observed dataset: are the values of a particular summary statistic computed from the simulated datasets reasonably close to the value of that statistic computed from the empirical data?

In phylogenetics, PPS has been largely limited to morphological or molecular character datasets (*e.g.*, Brown 2014; Höhna et al. 2018; Slater and Pennell 2014; May et al. 2021). This limited application of PPS is understandable, given that the character datasets are the only component of the study that is considered to be data under the standard phylogenetic model. However, for studies that rely on the tree model, such as diversification-rate analyses or divergence-time estimation, a more natural summary statistic would be one that relates to characteristics of the sample, rather than the morphological or molecular data. For example, if we wanted to assess the adequacy of a diversification model, we might use the number of samples at a particular time as a test statistic. The availability of this application of sample-based test statistics is one of the primary benefits—to theoreticians and empiricists alike—of recognizing the samples themselves as data that inform the tree model.

To demonstrate the utility of posterior-predictive distributions for samples under birth-death models, we applied this technique to the Marattiales analyses described in the Empirical Analyses section. We simulated datasets by simulating trees under the sampled fossilized birth-death model parameters, and keeping track of the number of fossils recorded in each geological epoch. The posterior-predictive distributions of the number of samples shows that the model with constant fossilizaton rates does a poor job of predicting the observed number of fossils in the Mississippian, Pennsylvanian, and Cisuralian (Fig. S7). By contrast, the model with variable fossilization rates does a much better job at predicting the number of fossils in these (and subsequent) intervals (Fig. S7, right). This result is concordant with our relative measures of model fit (using Bayes factors), which we report in the main text, and demonstrates that the variable-rate model is not only better-fitting than its constant-rate counterpart, but moreover that it is an *adequate representation* of the process that generated our data. (We present these results as a proof-of-concept rather than as a method: developing appropriate posterior-predictive methods is a significant task that requires validation and evaluation of statistic properties.)

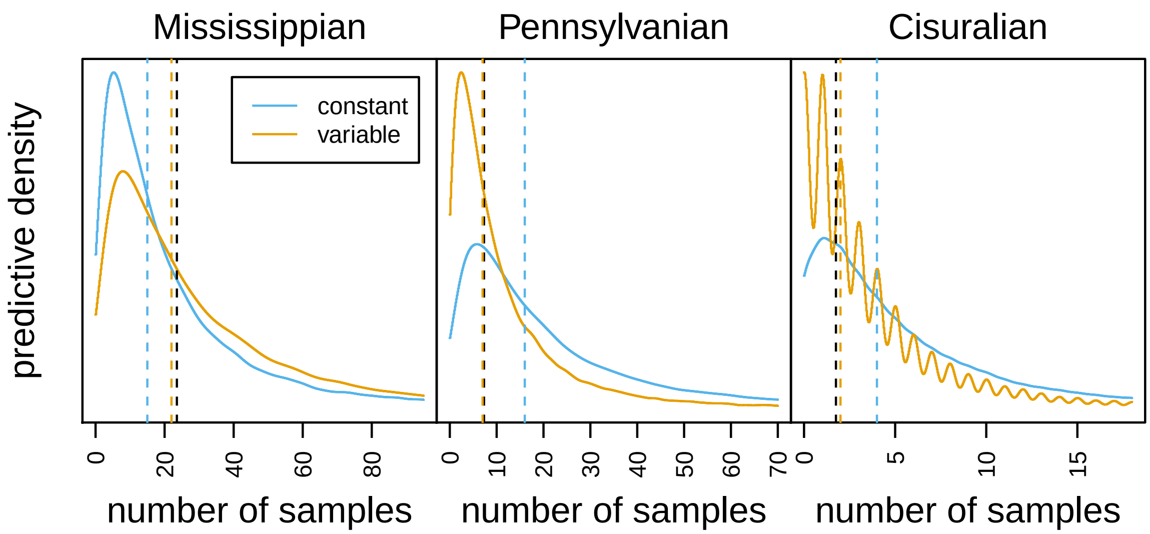


**Figure S7: Posterior-predictive simulation for the Marattiales dataset under two models.** We simulated fossil and extant marattialean datasets under a model with constant fossilization rates (blue) and variable fossilization rates (orange). Each density represents the posterior-predictive distribution for the number of samples in the given epoch. The black vertical line represent the observed number of samples in the epoch, and the colored vertical lines represent the median number of samples under the corresponding model.
